# TNF-α–Driven NOX5 Activation Promotes Oxidative Stress and Umbilical Artery Dysfunction in Preeclampsia

**DOI:** 10.64898/2026.05.18.726116

**Authors:** Nayara Carvalho-Barbosa, Mirele R. Machado, Juliano V. Alves, José Teles Oliveira-Neto, Josiane F. Silva, Ricardo C. Cavalli, Rita C. Tostes, Núbia S. Lobato, Rafael M. Costa

## Abstract

**Background:** Preeclampsia (PE) is a hypertensive disorder of pregnancy characterized by systemic inflammation, oxidative stress, and endothelial dysfunction. Although maternal vascular dysfunction is well established in PE, the mechanisms underlying fetal vascular injury remain poorly understood. We investigated whether inflammatory signaling activates NADPH oxidase 5 (NOX5) and contributes to oxidative stress and dysfunction in human umbilical arteries from pregnancies complicated by PE.

**Methods:** Umbilical arteries and serum samples were obtained from normotensive pregnant women (NP) and women with PE. Vascular reactivity, nitric oxide (NO) bioavailability, reactive oxygen species (ROS) generation, cytokine levels, and NOX isoform expression were evaluated in human umbilical arteries and EA.hy926 endothelial cells. Pharmacological inhibition of NOX5, TNF-α neutralization, Ca²⁺ channel blockade, and siRNA-mediated NOX5 silencing were used to investigate mechanisms.

**Results:** PE umbilical arteries exhibited increased vasoconstrictor responses, oxidative stress, and NOX5 expression, accompanied by impairment of NO bioavailability. NOX5 inhibition reversed vascular hyperreactivity in PE vessels. Exposure of normotensive umbilical arteries to PE serum reproduced the PE vascular phenotype, characterized by enhanced ROS generation, reduced NO levels, and hypercontractility. In endothelial cells, PE serum induced TNF-α-dependent Ca²⁺ influx, oxidative stress, and reduced NO production. Both pharmacological and genetic inhibition of NOX5 prevented these alterations.

**Conclusions:** PE promotes fetal vascular dysfunction through activation of a TNF-α/Ca^2+^/NOX5 signaling pathway that amplifies oxidative stress and impairs NO bioavailability. These findings identify NOX5 as a previously unrecognized mediator of umbilical artery dysfunction in PE and suggest the TNF-α/Ca^2+^/NOX5 axis as a potential therapeutic target in hypertensive pregnancies.

## INTRODUCTION

Preeclampsia (PE) is a multifactorial hypertensive disorder of pregnancy and a leading cause of maternal and fetal morbidity and mortality worldwide [1]. Although it affects approximately 5–8% of pregnancies, its clinical burden extends beyond gestation, predisposing both mother and offspring to long-term cardiovascular complications [2–4]. Despite extensive investigation, the mechanisms linking placental dysfunction to systemic and fetal vascular abnormalities remain incompletely understood.

PE is widely recognized as a two-stage disorder, initiated by defective placentation and followed by inadequate remodeling of the uterine spiral arteries, leading to placental ischemia and the release of bioactive factors into the maternal circulation [5–8]. These factors, including pro-inflammatory cytokines, anti-angiogenic mediators, and oxidative stress signals, drive widespread endothelial dysfunction and cardiovascular impairment [9–12]. While the maternal vascular consequences of PE have been extensively characterized, the impact of this hostile intrauterine environment on fetal vascular function, particularly in the umbilical circulation, remains less clearly defined.

Accumulating evidence indicates that oxidative stress is a central driver of PE pathophysiology. Increased production of reactive oxygen species (ROS), coupled with impaired antioxidant defenses, contributes to endothelial dysfunction, inflammation, and vascular dysregulation [13]. Among the enzymatic sources of ROS, NADPH oxidases have gained particular attention due to their regulated and stimulus-dependent activity [14]. However, the specific contribution of individual NADPH oxidase isoforms to fetal vascular dysfunction in PE remains a significant gap in knowledge.

NADPH oxidase 5 (NOX5) is a unique member of the NADPH oxidase family, distinguished by its direct activation through intracellular calcium (Ca²⁺) binding, independent of cytosolic regulatory subunits [15–17]. This feature positions NOX5 as a key transducer of Ca²⁺-dependent redox signaling, particularly under inflammatory and cellular stress conditions. Notably, NOX5 is expressed in endothelial and vascular smooth muscle cells, as well as in placental tissue, suggesting a potential role in regulating vascular tone and redox balance in both maternal and fetal compartments. Experimental models have demonstrated that NOX5 overexpression is associated with increased ROS production and elevated blood pressure, supporting its involvement in cardiovascular dysfunction [18–20].

In the context of PE, circulating pro-inflammatory cytokines such as tumor necrosis factor alpha (TNF-α) and interleukin 6 (IL-6) are markedly elevated and have been shown to disrupt Ca^2+^ homeostasis and endothelial function [21–25]. These alterations may provide a mechanistic link between systemic inflammation and NOX5 activation. However, whether this inflammatory-Ca**^2^**^+^ axis drives NOX5-dependent oxidative stress in the fetal vasculature, particularly in umbilical arteries, remains unknown.

Therefore, we hypothesized that, in PE, elevated circulating pro-inflammatory cytokines, especially the TNF-α, trigger intracellular Ca^2+^ accumulation in endothelial cells, thereby activating NOX5 and amplifying ROS generation. We further propose that this mechanism establishes a self-sustaining redox-inflammatory loop that impairs nitric oxide (NO) bioavailability and drives umbilical artery dysfunction.

## MATERIALS AND METHODS

The data that support the findings of this study are available from the corresponding author on reasonable request.

Detailed description of the methods used is available in the online-only Data Supplement.

## RESULTS

### Clinical characteristics of the study population

Clinical and biochemical characteristics of NP women and pregnant women with PE are summarized in Table 2. Women with PE exhibited significantly higher systolic and diastolic blood pressure, as well as marked proteinuria, confirming the hypertensive phenotype associated with the disease. In addition, circulating nitrite levels were significantly reduced in the PE group, suggesting impaired NO bioavailability. No significant differences were observed between groups regarding maternal age, gestational body mass index, or fasting glucose levels. Most women with PE received antihypertensive treatment with α-methyldopa, whereas a smaller proportion was treated with nifedipine. Consistent with the inflammatory profile associated with PE, circulating levels of TNF-α and IL-6 were increased in women with PE compared with NP controls, whereas IL-10 levels were reduced.

**Table 2.** Clinical and biochemical characteristics of NP women and women with PE.

| Parameters | NP<br>(n=12) | PE<br>(n=12) | p value |
| --- | --- | --- | --- |
| Gestational age at collection (weeks) | 41.0 ± 1.0 | 31.0 ± 2.0 | 0.0002 |
| Maternal age (years) | 28.0 ± 5.4 | 26.0 ± 5.9 | 0.8049 |
| Gestational BMI (kg/m<sup>2</sup>) | 22.8 ± 3.2 | 26.5 ± 4.7 | 0.5220 |
| SBP (mmHg) | 110.0 ± 8.5 | 140.0 ± 9.3 | 0.0263 |
| DBP (mmHg) | 70.0 ± 4.9 | 90.0 ± 5.8 | 0.0152 |
| Fasting glucose (mg/dL) | 74.0 ± 6.2 | 77.0 ± 6.1 | 0.7334 |
| Proteinuria (mg/24 h) | ND | 784.1 ± 205.3 | 0.0009 |
| Plasma nitrite (nM) | 147.2 ± 49.4 | 79.1 ± 23.8 | 0.0477 |
| TNF-α (pg/mL) | 5.3 ± 0.7 | 19.8 ± 0.9 | 0.0001 |
| IL-6 (pg/mL) | 4.2 ± 0.6 | 16.3 ± 1.1 | 0.0001 |
| IL-10 (pg/mL) | 44.7 ± 1.2 | 14.1 ± 1.6 | 0.0004 |
| **Antihypertensive drugs** |  |  |  |
| α-Methyldopa (%) | NA | 75.0 | |
| Nifedipine (%) | NA | 25.0 |  |

Values are presented as mean ± SEM (n=12). BMI, body mass index; SBP, systolic blood pressure; DBP, diastolic blood pressure; TNF-α , tumor necrosis factor alpha; IL-6, interleukin 6; IL-10, interleukin 10; ND, not detectable; NA, not applicable. Student's t-test: p < 0.05 vs. normotensive pregnant women (NP).

### Umbilical arteries from pregnancies with PE exhibit enhanced contractile responses and reduced NO bioavailability

To investigate whether PE is associated with functional alterations in the fetal vasculature, vascular reactivity was evaluated in human umbilical arteries obtained from NP women and women with PE. Endothelium-independent relaxation induced by sodium nitroprusside was similar between groups, indicating preserved vascular smooth muscle responsiveness to exogenous NO in PE umbilical arteries (Figure 1A). In contrast, contractile responses were significantly augmented in umbilical arteries from PE pregnancies. Stimulation with the thromboxane A₂ analog U46619 produced greater vasoconstriction in PE vessels compared with NP controls (Figure 1B). Likewise, serotonin-induced contractions were markedly increased in arteries obtained from PE pregnancies (Figure 1C), suggesting an overall hypercontractile phenotype in the fetal vasculature during PE. To determine whether impaired endogenous NO signaling contributed to this enhanced vasoconstriction, vascular responses to serotonin were evaluated in the presence of the NOS inhibitor L-NAME. In umbilical arteries from NP pregnancies, NOS inhibition significantly potentiated serotonin-induced contraction, indicating an important modulatory role of endogenous NO under physiological conditions. In contrast, L-NAME failed to further increase contractile responses in PE arteries (Figure 1D), suggesting that endogenous NO-mediated vascular modulation is already compromised in PE. Consistent with this interpretation, nitrite levels were significantly reduced in umbilical arteries from PE pregnancies compared with NP controls (Figure 1E). Moreover, direct assessment of NO production using DAF-2 DA fluorescence demonstrated markedly lower NO levels in PE umbilical arteries (Figure 1F). Together, these findings demonstrate that PE is associated with impaired NO bioavailability and increased vasoconstrictor responsiveness in the umbilical circulation.

**Figure 1.**
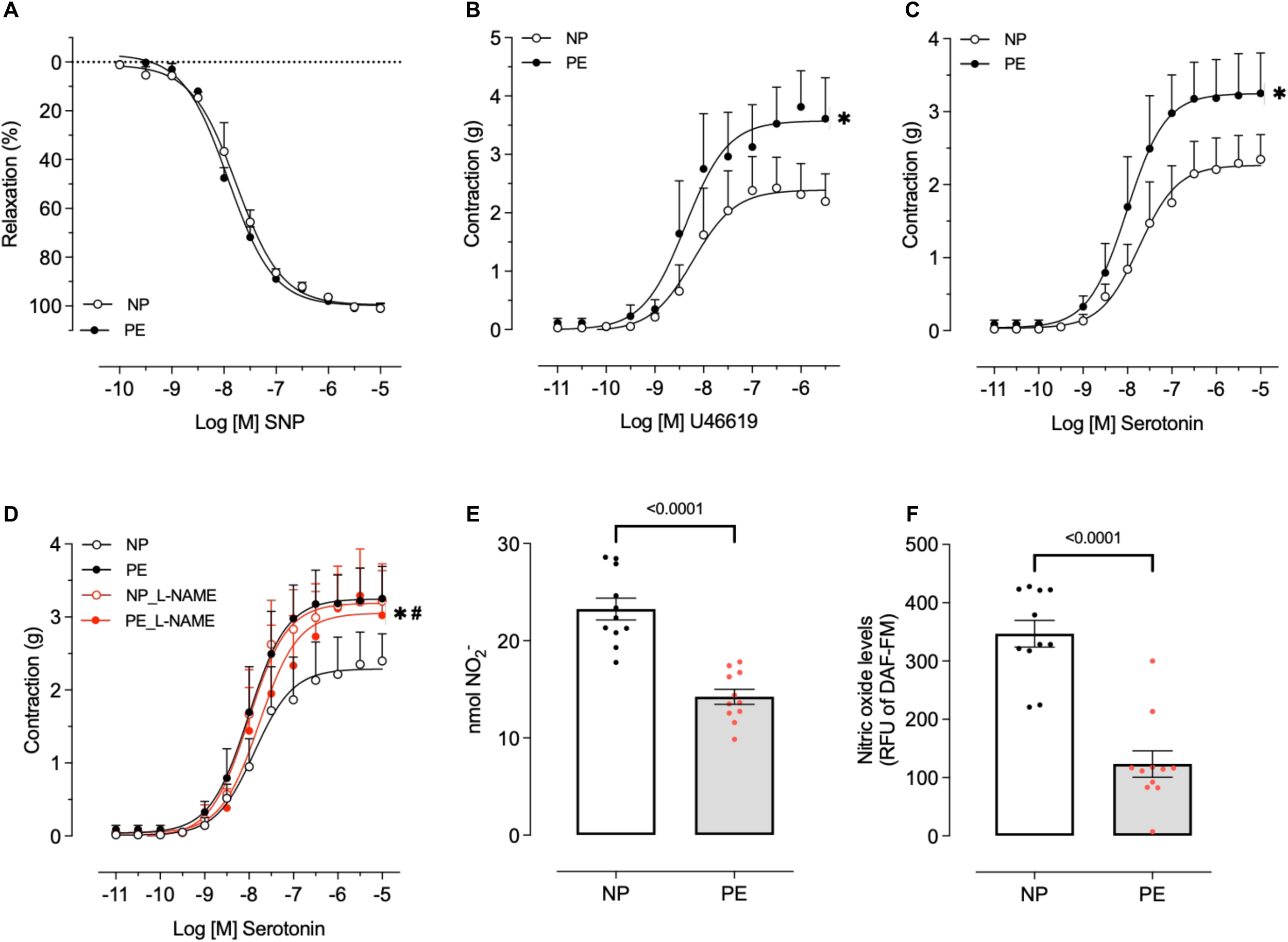
Reduced nitric oxide bioavailability contributes to enhanced serotonin-induced vascular reactivity in umbilical cord arteries from preeclamptic pregnancies. Concentration–effect curves for sodium nitroprusside (A), the thromboxane A₂ analog U46619 (B), and serotonin (C) were obtained in umbilical cord arteries from normotensive (NP) and preeclamptic (PE) pregnant women. Additional concentration–effect curves for serotonin were performed in the presence of vehicle or the nitric oxide synthase inhibitor L-NAME (100 μM, 30 min). Nitric oxide bioavailability was assessed using the Griess assay (E) and diaminofluorescein-FM (DAF-FM) fluorescence (F). Values are expressed as mean ± SEM (n = 11-12). Statistical analysis was performed using ANOVA or Student’s t-test.

### Increased oxidative stress and NOX5 upregulation contribute to vascular dysfunction in umbilical arteries from pregnancies with PE

Given the marked impairment in NO bioavailability and enhanced vasoconstrictor responses observed in PE umbilical arteries, we next investigated whether oxidative stress contributes to this dysfunctional vascular phenotype. DHE fluorescence revealed a significant increase in ROS production in umbilical arteries obtained from PE pregnancies compared with NP controls (Figure 2A). Consistently, lucigenin-enhanced chemiluminescence demonstrated increased NADPH-dependent ROS generation in PE vessels (Figure 2B). In addition, H₂O₂ production was significantly elevated in umbilical arteries from PE pregnancies (Figure 2C), further supporting the presence of an exacerbated oxidative environment in the fetal vasculature during PE. Because NOX5 is a Ca²⁺-sensitive NADPH oxidase isoform strongly associated with vascular redox signaling, we next evaluated its expression in umbilical arteries. PE vessels exhibited significantly increased NOX5 mRNA expression compared with NP arteries (Figure 2D). Similarly, protein content analysis showed an increase in NOX5 content in the umbilical arteries of pregnancies with PE (Figure 2E). To determine whether NOX5 contributes functionally to the hypercontractile phenotype observed in PE arteries, vascular responses to serotonin were assessed in the presence of melittin, a NOX5 inhibitor. Melittin did not affect serotonin-induced contraction in umbilical arteries from NP pregnancies. In contrast, pharmacological inhibition of NOX5 markedly reduced the exaggerated contractile response observed in PE arteries, restoring vascular reactivity to levels comparable to those observed in NP vessels (Figure 2F). These findings suggest that increased NOX5-derived oxidative stress is a major contributor to umbilical artery dysfunction in PE.

**Figure 2.**
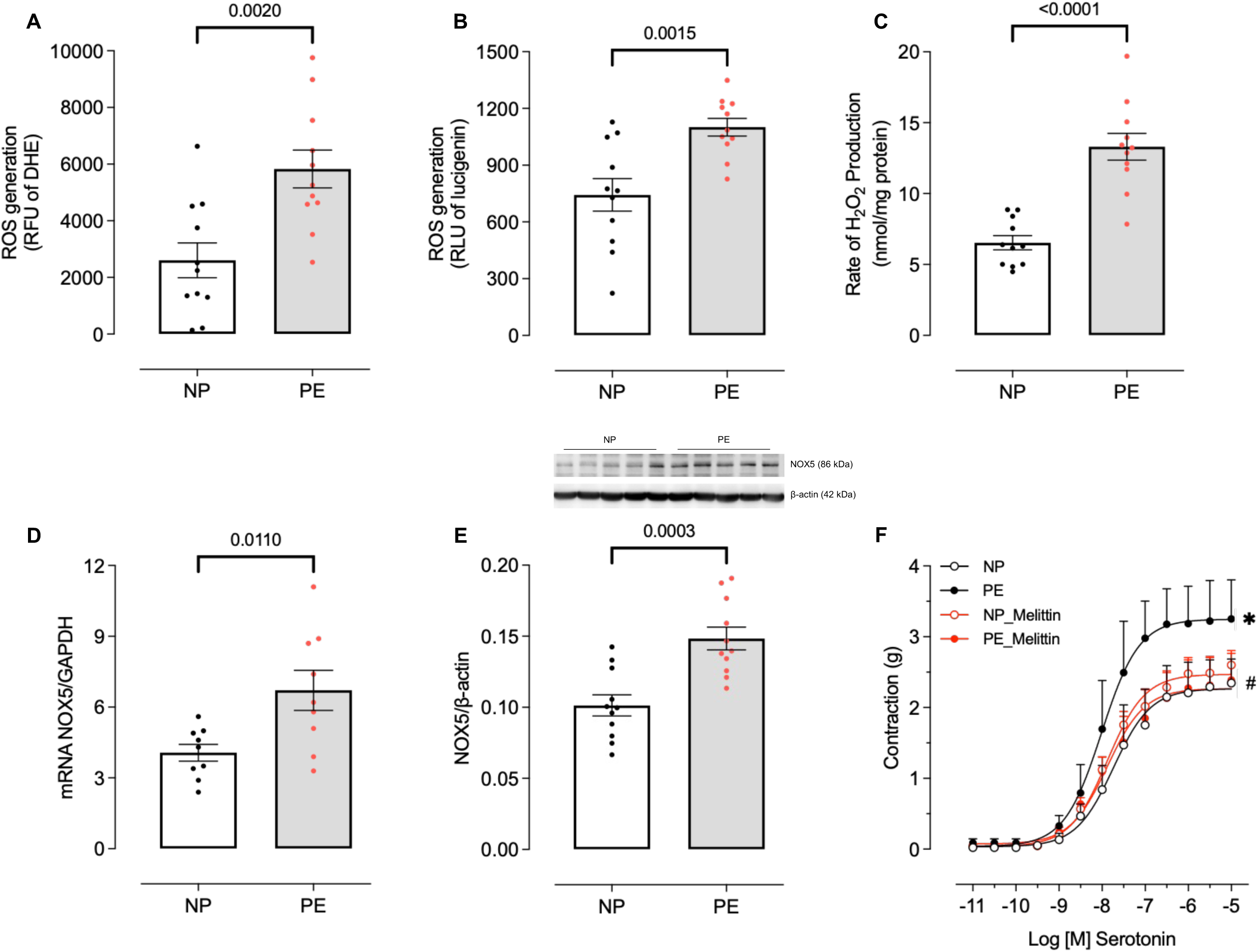
Increased oxidative stress and NOX5 upregulation contribute to enhanced vascular reactivity in umbilical cord arteries from preeclamptic pregnancies. Reactive oxygen species (ROS) production was evaluated in umbilical cord arteries from normotensive (NP) and preeclamptic (PE) pregnant women using dihydroethidium (DHE) fluorescence (A) and lucigenin-enhanced chemiluminescence (B). Hydrogen peroxide (H₂O₂) levels were measured using the Amplex Red assay (C). NOX5 expression was assessed at the mRNA (D) and protein levels (E). Concentration–effect curves for serotonin were performed in umbilical cord arteries from NP and PE groups in the presence of vehicle or the NOX5 inhibitor melittin (0.1 μM, 30 min) (F). Values are expressed as mean ± SEM (n = 11-12). Statistical analysis was performed using ANOVA or Student’s t-test.

Importantly, the increase in NOX5 expression appeared to be selective, since no differences were observed in NOX1 or NOX4 mRNA expression between groups (Supplementary Figure 1A–B). Likewise, protein content of NOX1 and NOX4 was similar in NP and PE arteries (Supplementary Figure 1C–D), supporting a predominant involvement of NOX5 in the oxidative phenotype associated with PE. We next investigated whether alterations in antioxidant defense systems could additionally contribute to ROS accumulation in PE arteries. No differences were observed in SOD1 protein content or enzymatic activity between groups (Supplementary Figure 2A and 2C). In contrast, catalase protein content and enzymatic activity were both significantly reduced in PE umbilical arteries compared with NP controls (Supplementary Figure 2B and 2D). These findings suggest that reduced catalase activity may contribute to H₂O₂ accumulation in PE umbilical arteries.

### Circulating factors from women with PE induce NOX5-dependent oxidative stress and vascular dysfunction in normotensive umbilical arteries

To determine whether circulating factors present in PE contribute to NOX5 activation and vascular dysfunction, umbilical arteries obtained from NP pregnancies were incubated with serum from NP or PE women. Incubation with PE serum significantly increased ROS production in NP umbilical arteries, as demonstrated by increased DHE fluorescence (Figure 3A), enhanced lucigenin chemiluminescence (Figure 3B), and elevated H₂O₂ production (Figure 3C). Importantly, these effects were prevented by incubation with melittin, indicating that PE serum induces NOX5-dependent oxidative stress in otherwise healthy umbilical arteries. We next evaluated whether exposure to PE serum also affects NO bioavailability. Incubation of NP umbilical arteries with PE serum significantly reduced nitrite levels (Figure 3D) and decreased NO production assessed by DAF-2 DA fluorescence (Figure 3E). Notably, pharmacological inhibition of NOX5 prevented the reduction in NO bioavailability induced by PE serum, further supporting a mechanistic link between circulating PE-associated factors, NOX5 activation, and oxidative stress-mediated NO depletion. To investigate the functional consequences of these alterations, vascular reactivity to serotonin was evaluated in NP umbilical arteries incubated with NP or PE serum. Exposure to PE serum markedly increased serotonin-induced contraction compared with arteries incubated with NP serum (Figure 3F). This hypercontractile response was prevented by melittin, whereas incubation with NP serum did not affect vascular responsiveness. Together, these findings demonstrate that circulating factors present in PE promote NOX5-dependent oxidative stress, impair NO bioavailability, and induce vascular hyperreactivity in human umbilical arteries.

**Figure 3.**
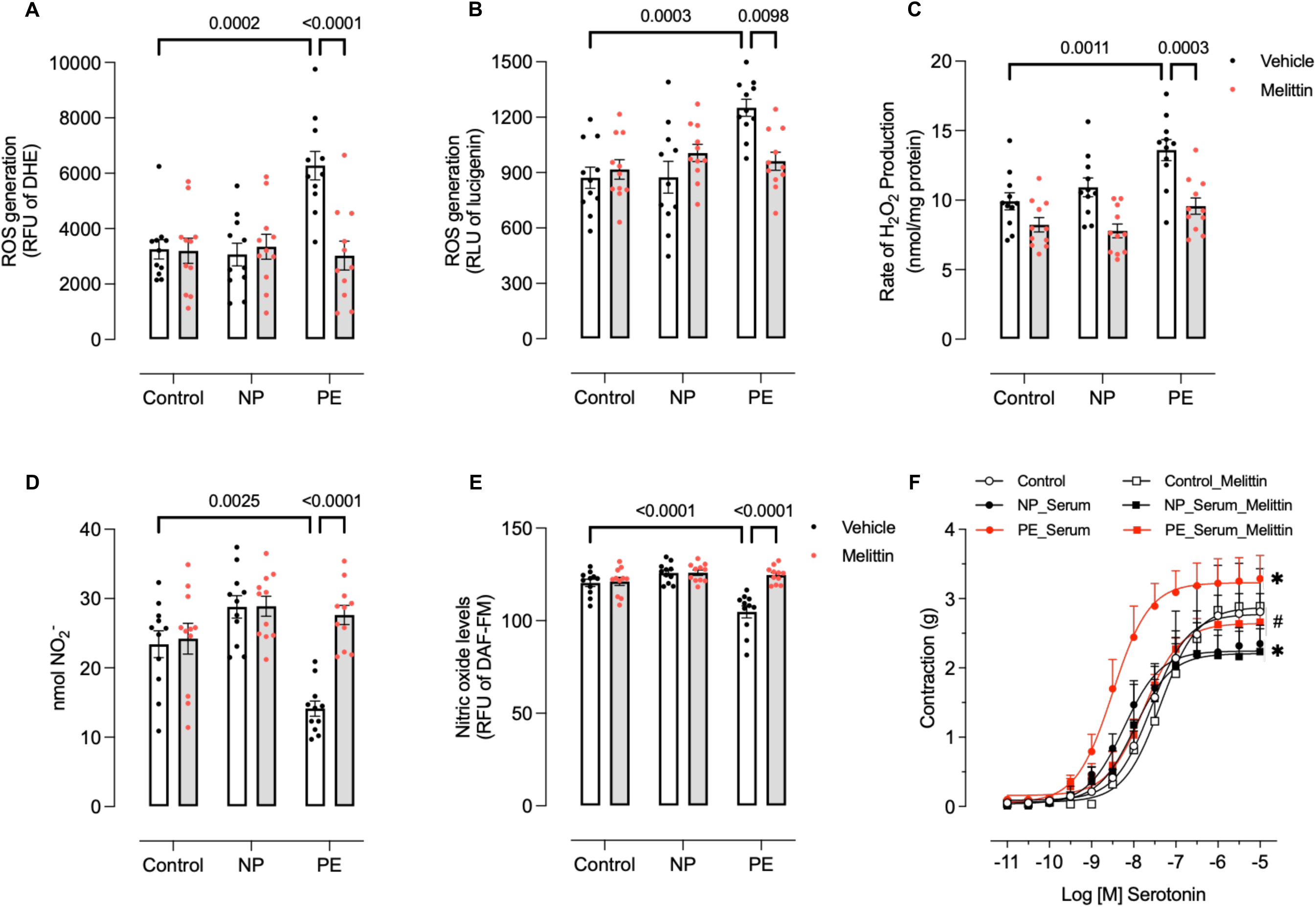
Preeclamptic serum induces oxidative stress, reduces nitric oxide bioavailability, and enhances vascular reactivity in umbilical cord arteries. Umbilical cord arteries from normotensive (NP) pregnant women were incubated with fetal bovine serum (FBS, control) or serum from normotensive (NP) or preeclamptic (PE) pregnant women (20% v/v, 1 h). Reactive oxygen species (ROS) production was assessed by dihydroethidium (DHE) fluorescence (A) and lucigenin-enhanced chemiluminescence (B). Hydrogen peroxide (H₂O₂) levels were measured using the Amplex Red assay (C). Nitric oxide (NO) bioavailability was assessed using the Griess assay (D) and diaminofluorescein-FM (DAF-FM) fluorescence (E). Concentration–effect curves to serotonin were obtained in umbilical cord arteries incubated with FBS, NP serum, or PE serum, in the presence of vehicle or melittin (0.1 μM, 30 min) (F). Values are expressed as mean ± SEM (n = 11-12). Statistical analysis was performed using ANOVA.

### Circulating factors from women with PE promote TNF-α-dependent calcium influx in endothelial cells

To investigate the cellular mechanisms underlying NOX5 activation in PE, experiments were performed in EA.hy926 endothelial cells stimulated with serum obtained from NP or PE women. Initially, cell viability was assessed to determine whether serum stimulation affected endothelial cell integrity. Exposure to NP or PE serum (20% v/v for 1 h) did not alter cell viability under the experimental conditions employed (Supplementary Figure 3). Because NOX5 activity is tightly regulated by intracellular Ca²⁺ levels, we next evaluated calcium influx in endothelial cells following serum stimulation. Incubation with NP serum did not significantly affect intracellular Ca²⁺ levels, whereas stimulation with PE serum markedly increased Ca²⁺ influx (Figure 4A). Quantitative analysis of the area under the curve (AUC) confirmed the significant increase in intracellular Ca²⁺ mobilization induced by PE serum (Figure 4B). Considering the elevated circulating TNF-α levels observed in women with PE, we next investigated whether TNF-α directly contributes to endothelial Ca²⁺ influx. Stimulation with recombinant TNF-α induced a concentration-dependent increase in intracellular Ca²⁺ levels, with greater responses observed at higher cytokine concentrations (Figure 4C). These effects were confirmed by AUC analysis (Figure 4D), supporting a direct role for TNF-α in endothelial Ca²⁺ mobilization. To determine whether TNF-α contributes to the effects induced by PE serum, endothelial cells were incubated with infliximab, a TNF-α-neutralizing antibody. Infliximab did not alter intracellular Ca²⁺ levels in cells stimulated with NP serum. However, TNF-α neutralization partially attenuated the increase in Ca²⁺ influx induced by PE serum (Figure 4E–F), suggesting that TNF-α is an important, although not exclusive, circulating mediator involved in this response. Finally, to investigate the involvement of Ca²⁺ channels in PE serum-induced Ca²⁺ influx, cells were incubated with nifedipine, an L-type Ca²⁺ channel blocker. Nifedipine had no effect in cells stimulated with NP serum but markedly inhibited the increase in intracellular Ca²⁺ induced by PE serum (Figure 4G–H). Together, these findings demonstrate that circulating factors present in PE, particularly TNF-α, promote endothelial Ca²⁺ influx through activation of Ca²⁺ channels, providing a mechanistic basis for NOX5 activation in PE.

**Figure 4.**
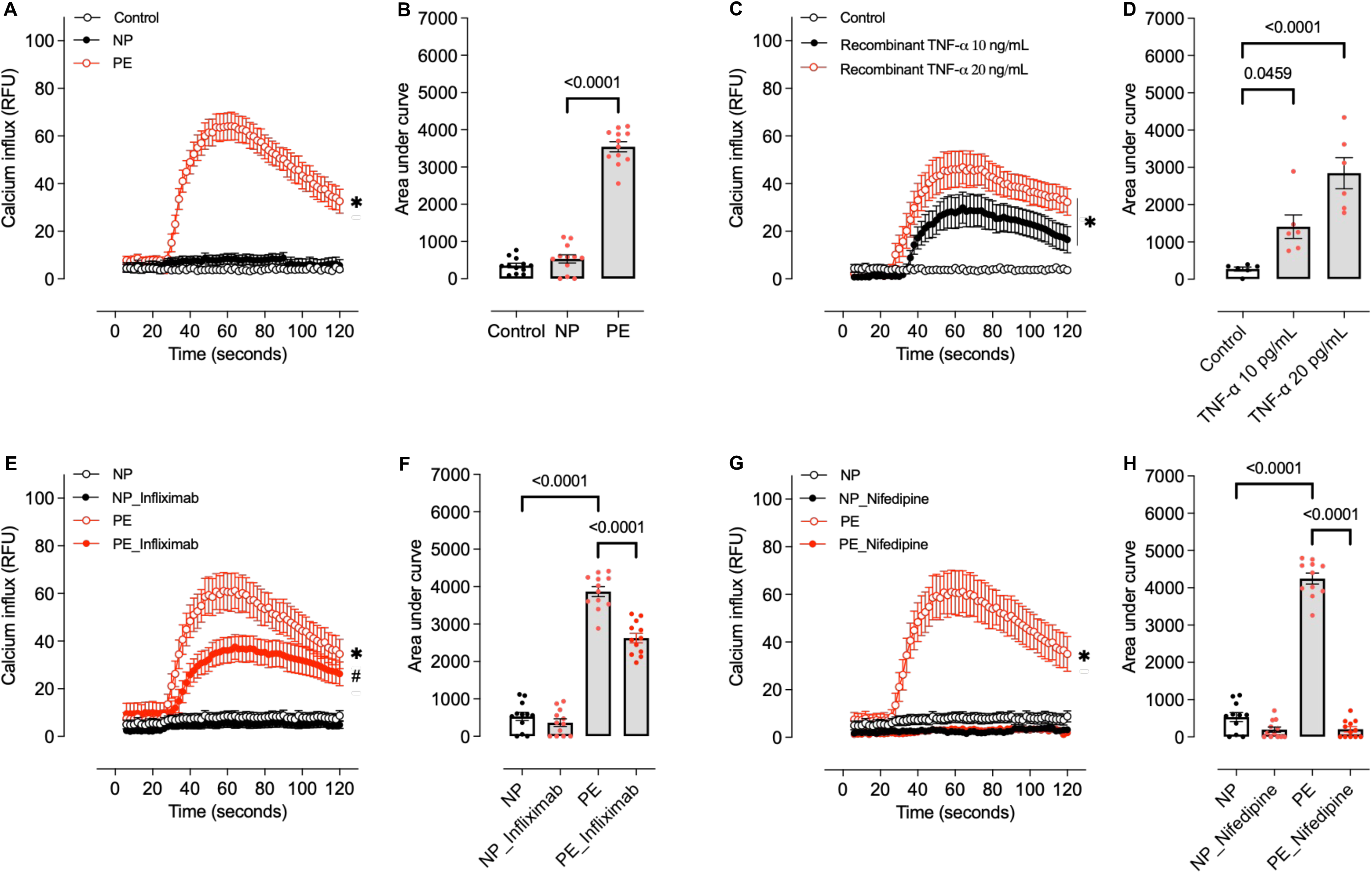
TNF-α–dependent calcium influx is enhanced by preeclamptic serum in endothelial cells. Intracellular Ca²⁺ influx was measured as relative fluorescence units (RFU) in EA.hy926 endothelial cells following stimulation with recombinant TNF-α or incubation with serum from normotensive (NP) or preeclamptic (PE) pregnant women (20% v/v, 1 h). Representative fluorescence traces and the corresponding area under the curve (AUC) are shown under basal conditions (A, B), after stimulation with recombinant TNF-α (C, D), and following pre-treatment with the anti–TNF-α antibody infliximab (1 μM, 30 min; E, F) or the L-type calcium channel blocker nifedipine (1 μM, 30 min; G, H). Values are expressed as mean ± SEM (n = 6-12). Statistical analysis was performed using ANOVA.

### TNF-α, Ca^2+^ influx, and NOX5 activation mediate oxidative stress and NO depletion induced by preeclamptic serum in endothelial cells

We next investigated whether circulating factors present in PE serum directly impair endothelial NO bioavailability and promote oxidative stress in EA.hy926 endothelial cells. Stimulation with PE serum significantly reduced nitrite levels compared with cells exposed to NP serum (Figure 5A). Similarly, direct NO measurement using DAF-2 DA fluorescence demonstrated a marked reduction in NO production following stimulation with PE serum (Figure 5B). In parallel, endothelial cells exposed to PE serum exhibited increased ROS generation, as demonstrated by enhanced lucigenin chemiluminescence (Figure 5C) and increased H_2_O_2_ production (Figure 5D), confirming that circulating factors present in PE induce a robust pro-oxidative endothelial phenotype. To further investigate whether PE serum modulates NADPH oxidase isoforms in endothelial cells, we evaluated the gene expression of NOX1, NOX4, and NOX5. Exposure to PE serum significantly increased NOX1 and NOX5 mRNA expression (Supplementary Figure 4A and 4C), whereas NOX4 expression remained unchanged (Supplementary Figure 4B). Importantly, TNF-α neutralization with infliximab prevented the increase in NOX5 expression induced by PE serum, suggesting that TNF-α signaling contributes to NOX5 upregulation in endothelial cells. To investigate the mechanisms involved in these responses, endothelial cells were incubated with infliximab, nifedipine, or melittin prior to PE serum stimulation. TNF-α neutralization with infliximab, blockade of L-type Ca²⁺ channels with nifedipine, and NOX5 inhibition with melittin all prevented the reduction in nitrite levels and NO production induced by PE serum (Figure 5A–B). Likewise, all three interventions markedly attenuated ROS and H₂O₂ generation triggered by PE serum exposure (Figure 5C–D). Together, these findings demonstrate that circulating factors present in PE serum promote endothelial oxidative stress and NO depletion through a mechanism involving TNF-α signaling, Ca²⁺ influx, and NOX5 activation.

**Figure 5.**
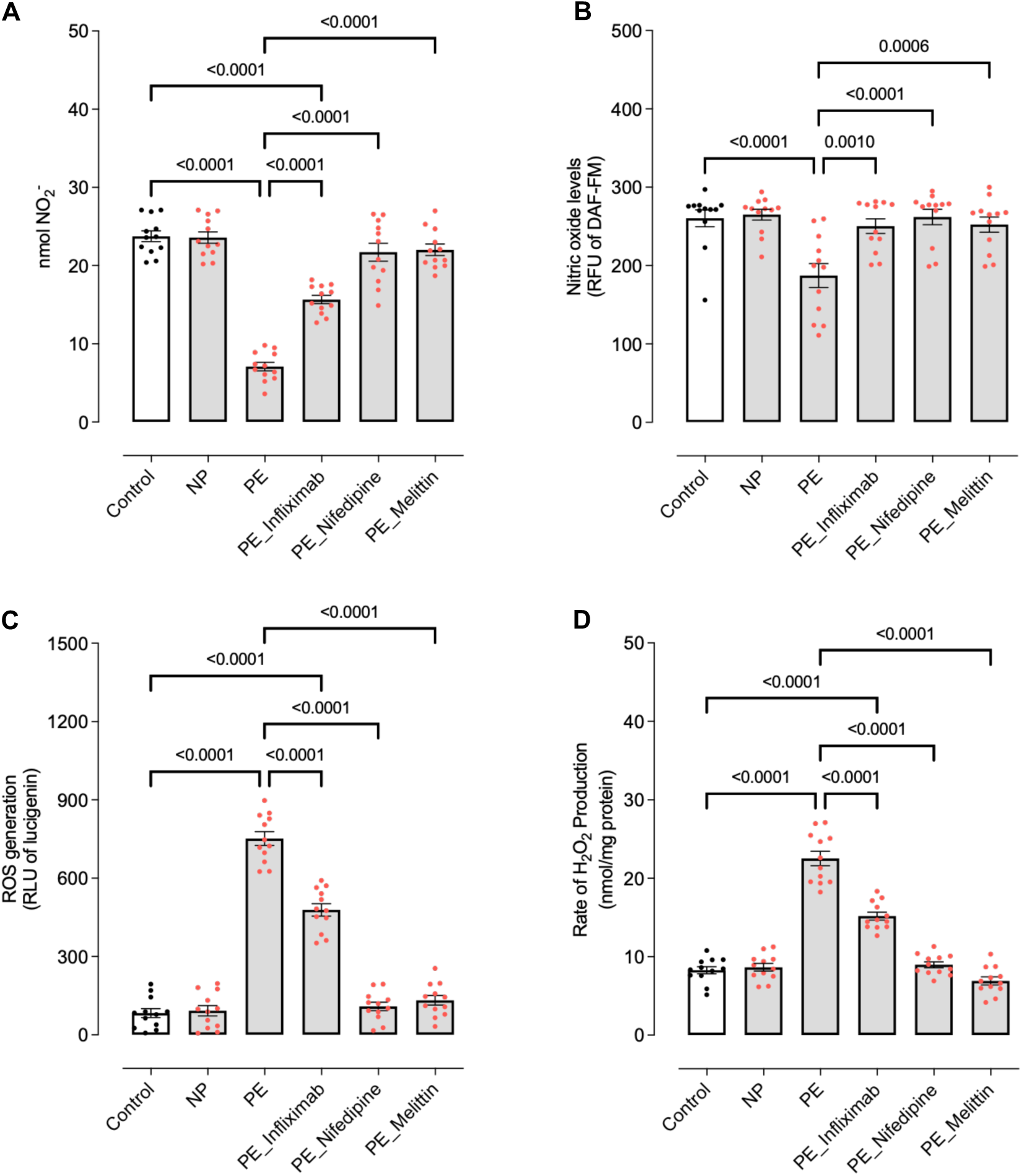
Preeclamptic serum reduces nitric oxide bioavailability and increases oxidative stress in endothelial cells via TNF-α- and Ca²⁺-dependent mechanisms. EA.hy926 endothelial cells were incubated with fetal bovine serum (FBS, control) or serum from normotensive (NP) or preeclamptic (PE) pregnant women (20% v/v, 1 h). Cells were treated with vehicle or the anti–TNF-α antibody infliximab (1 μM, 30 min), the L-type calcium channel blocker nifedipine (1 μM, 30 min), or the NOX5 inhibitor melittin (0.1 μM, 30 min). Nitric oxide (NO) bioavailability was assessed using the Griess assay (A) and diaminofluorescein-FM (DAF-FM) fluorescence (B). Reactive oxygen species (ROS) production was measured by lucigenin-enhanced chemiluminescence (C), and hydrogen peroxide (H_2_O_2_) levels were determined using the Amplex Red assay (D). Values are expressed as mean ± SEM (n = 11-12). Statistical analysis was performed using ANOVA.

### Genetic silencing of NOX5 prevents oxidative stress and NO depletion induced by preeclamptic serum in endothelial cells

To further establish the role of NOX5 in the endothelial dysfunction induced by PE serum, EA.hy926 endothelial cells were subjected to NOX5 gene silencing using siRNA-mediated knockdown. Western blot analysis confirmed that NOX5 siRNA markedly reduced NOX5 protein content compared with control conditions (Figure 6A). Consistent with previous findings, stimulation of endothelial cells with PE serum significantly reduced nitrite levels (Figure 6B) and NO production assessed by DAF-2 DA fluorescence (Figure 6C), while simultaneously increasing ROS generation measured by lucigenin chemiluminescence (Figure 6D) and H_2_O_2_ production (Figure 6E). Importantly, NOX5 silencing prevented the reduction in nitrite levels and NO bioavailability induced by PE serum. Likewise, siRNA-mediated NOX5 knockdown markedly attenuated ROS and H₂O₂ generation triggered by PE serum exposure. These findings demonstrate that NOX5 is a critical mediator of the pro-oxidative and NO-depleting effects induced by circulating factors present in PE serum. Together, these data provide genetic evidence supporting the pharmacological findings obtained with melittin and further establish NOX5 as a central mediator of endothelial oxidative stress and dysfunction associated with PE.

**Figure 6.**
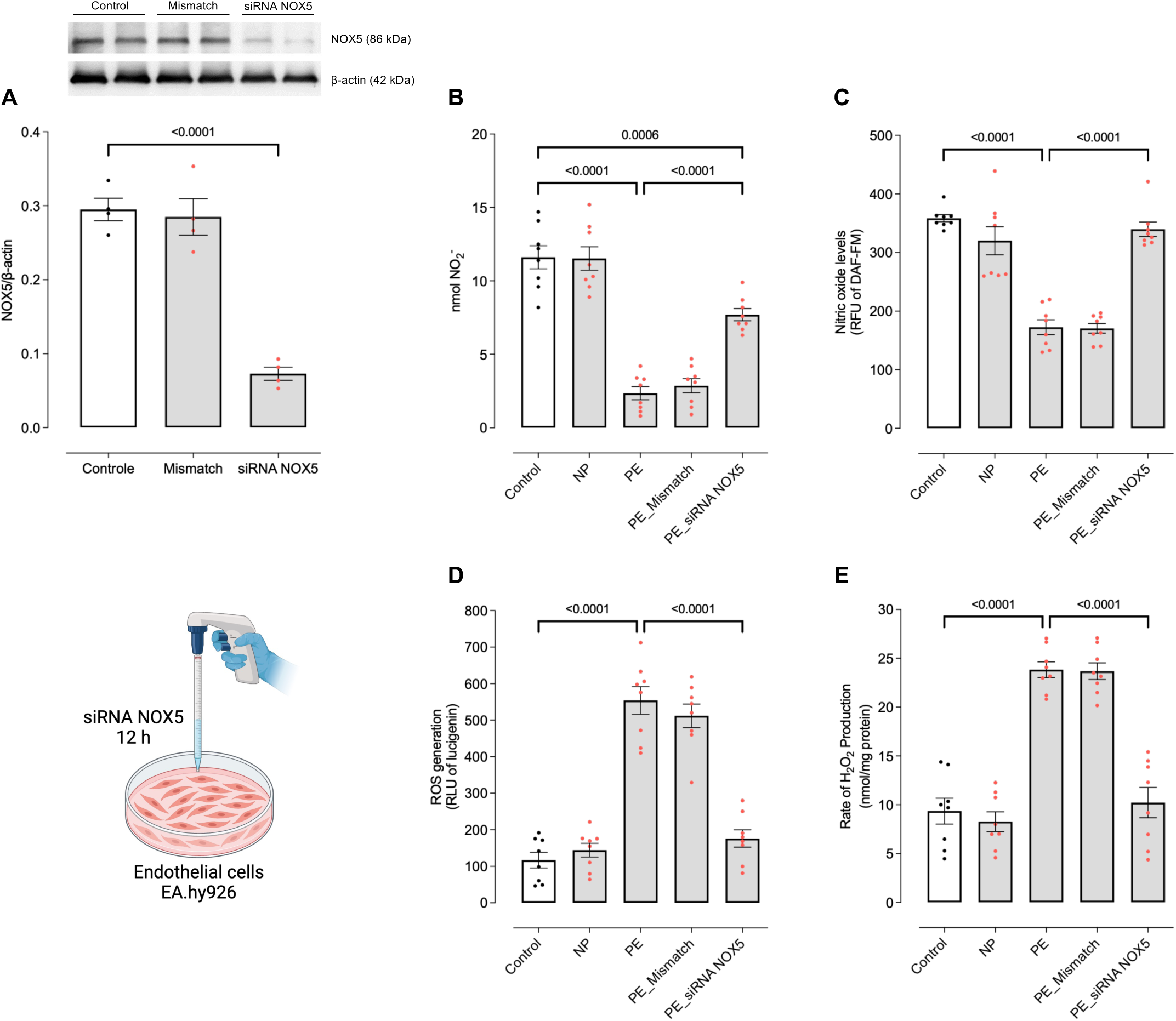
NOX5 silencing prevents preeclamptic serum–induced oxidative stress and restores nitric oxide bioavailability in endothelial cells. EA.hy926 endothelial cells were transfected with NOX5 siRNA for 12 h to achieve gene silencing, which was confirmed by protein expression analysis (Western blot; A). Following transfection, cells were incubated with fetal bovine serum (FBS, control) or serum from normotensive (NP) or preeclamptic (PE) pregnant women (20% v/v, 1 h). Nitric oxide (NO) bioavailability was assessed using the Griess assay (B) and diaminofluorescein-FM (DAF-FM) fluorescence (C). Reactive oxygen species (ROS) production was measured by lucigenin-enhanced chemiluminescence (D), and hydrogen peroxide (H₂O₂) levels were determined using the Amplex Red assay (E). Values are expressed as mean ± SEM (n = 4-8). Statistical analysis was performed using ANOVA.

## DISCUSSION

The present study identifies NOX5 as a previously unrecognized mediator of fetal vascular dysfunction in PE and provides mechanistic evidence linking systemic inflammation to endothelial oxidative stress through a TNF-α/Ca²⁺-dependent pathway. Although oxidative stress and endothelial dysfunction are well-established features of PE, the enzymatic sources responsible for ROS generation within the fetal vasculature remain incompletely understood. Here, we demonstrate that umbilical arteries from PE pregnancies exhibit marked hypercontractility associated with increased ROS production, reduced NO bioavailability, and selective upregulation of NOX5. Importantly, both pharmacological inhibition and siRNA-mediated silencing of NOX5 prevented the oxidative and nitrosative imbalance induced by PE serum, establishing NOX5 not merely as a marker of oxidative stress, but as a functional driver of endothelial dysfunction in PE.

Previous studies have largely focused on placental oxidative stress and maternal endothelial dysfunction [26,27], whereas comparatively little attention has been directed toward the fetal vasculature. This is particularly relevant because abnormal umbilical artery Doppler indices are strongly associated with fetal growth restriction, placental insufficiency, and adverse perinatal outcomes in pregnancies complicated by PE [28–30]. Our findings therefore expand the current paradigm by demonstrating that PE-associated vascular injury extends directly to the umbilical circulation. Moreover, the observation that PE serum reproduced vascular hyperreactivity and oxidative stress in normotensive arteries indicates that circulating pathogenic mediators are sufficient to transfer the dysfunctional phenotype, supporting the concept that PE behaves as a systemic inflammatory vascular disorder rather than a disease restricted to placental pathology.

Among NADPH oxidase isoforms, NOX5 is uniquely positioned to participate in PE-associated vascular dysfunction because its activation is directly dependent on intracellular Ca²⁺. Unlike NOX1 and NOX4, which were unchanged in our study, NOX5 contains EF-hand Ca²⁺-binding domains that couple inflammatory signaling to rapid ROS generation [31–33]. This characteristic becomes particularly relevant in the context of PE, a condition marked by cytokine excess and dysregulated endothelial Ca²⁺ handling. Indeed, PE serum and recombinant TNF-α both increased endothelial Ca²⁺ influx, whereas TNF-α neutralization and Ca²⁺ channel blockade attenuated NOX5-associated oxidative stress. These findings support a mechanistic model in which inflammatory mediators activate endothelial Ca²⁺ signaling upstream of NOX5, thereby amplifying ROS generation and impairing NO bioavailability.

The translational relevance of these observations is strengthened by the fact that NOX5 is absent in rodents, which may partially explain why its contribution to PE has remained underexplored despite increasing evidence implicating NOX5 in human cardiovascular disease [34]. The use of human umbilical arteries and human endothelial cells therefore represents an important strength of the present study, allowing investigation of a clinically relevant redox pathway that cannot be adequately reproduced in conventional murine models.

Another important finding was the reduction in catalase expression and activity in PE arteries. Because catalase is a major determinant of H_2_O_2_ detoxification, impaired antioxidant defense may favor sustained H_2_O_2_ accumulation within the vascular wall [35]. Beyond its oxidative effects, H_2_O_2_ acts as a redox-sensitive signaling molecule capable of modulating vascular contraction, endothelial permeability, and inflammatory pathways [36,37]. Thus, the coexistence of increased NOX5-derived ROS generation and impaired antioxidant buffering suggests the establishment of a self-sustaining oxidative microenvironment within the fetal vasculature during PE.

From a clinical perspective, our findings may also have implications for current antihypertensive strategies used in PE. Nifedipine is widely employed for blood pressure control during hypertensive pregnancies [38,39]; however, our data suggest that blockade of Ca²⁺ influx additionally limits NOX5 activation and endothelial oxidative stress. Although speculative, this raises the possibility that part of the vascular benefit of nifedipine may extend beyond hemodynamic control and involve modulation of redox-sensitive pathways.

Despite the strengths of the present study, some limitations should be acknowledged. First, the relatively small sample size reflects the inherent difficulty in obtaining and processing fresh human umbilical artery samples immediately after delivery, particularly from pregnancies complicated by early-onset PE. Nevertheless, the consistency observed across vascular, molecular, biochemical, pharmacological, and gene-silencing approaches substantially strengthens the robustness of the findings. Second, most patients included in the PE group presented early-onset disease and were receiving antihypertensive therapy, mainly α-methyldopa and nifedipine, which may have partially influenced vascular and inflammatory parameters. However, significant endothelial dysfunction and oxidative stress were still observed despite ongoing treatment, underscoring the severity of the vascular phenotype associated with PE. Third, although EA.hy926 cells represent a well-established endothelial model, they do not fully recapitulate the complexity of primary fetoplacental endothelial cells in vivo. Finally, while melittin was used as a pharmacological NOX5 inhibitor, this compound may exert biological effects beyond NOX5 inhibition [40,41]. Importantly, however, the use of siRNA-mediated NOX5 silencing provided complementary genetic evidence supporting the central role of NOX5 in the mechanisms described herein.

In conclusion, the present study identifies NOX5 as a key mediator of fetal vascular dysfunction in PE and uncovers a mechanistic pathway linking systemic inflammation to endothelial oxidative stress through TNF-α-dependent Ca²⁺ signaling. Our findings demonstrate that circulating factors present in PE promote NOX5 activation, excessive ROS generation, impaired NO bioavailability, and vascular hyperreactivity in the umbilical circulation. By integrating human vascular physiology, redox biology, inflammatory signaling, pharmacological interventions, and gene-silencing approaches, this study advances current understanding of the mechanisms underlying endothelial dysfunction in PE and positions the TNF-α/Ca²⁺/NOX5 axis as a potential therapeutic target in hypertensive disorders of pregnancy. More broadly, these findings reinforce the concept that PE is not only a maternal vascular disease, but also a condition capable of directly programming fetal vascular injury through redox-dependent mechanisms.

## PERSPECTIVE

The present study identifies NOX5 as a previously unrecognized mediator of fetal vascular dysfunction in PE and highlights the TNF-α/Ca²⁺/NOX5 axis as a potential therapeutic target in hypertensive pregnancies. Beyond advancing the understanding of redox-dependent mechanisms in the umbilical circulation, these findings suggest that modulation of NOX5 activity may represent a novel strategy to preserve fetal vascular homeostasis and reduce adverse perinatal outcomes associated with PE. Future studies are warranted to investigate the long-term cardiovascular implications of NOX5-driven fetal vascular injury and to evaluate the therapeutic potential of targeting this pathway in clinical settings.

## NONSTANDARD ABBREVIATIONS AND ACRONYMS

AUC: Area under the curve
BMI: Body mass index
Ca²⁺: Calcium
DAF-2 DA: 4,5-Diaminofluorescein diacetate
DBP: Diastolic blood pressure
DHE: Dihydroethidium
DMEM: Dulbecco’s Modified Eagle Medium
DMSO: Dimethyl sulfoxide
EA.hy926: Human endothelial cell line EA.hy926
EC_50_: Half-maximal effective concentration
ELISA: Enzyme-linked immunosorbent assay
Emax: Maximal response
FBS: Fetal bovine serum
HBSS: Hanks’ balanced salt solution
HRP: Horseradish peroxidase
H_2_O_2_: Hydrogen peroxide
IL-6: Interleukin 6
IL-10: Interleukin 10
L-NAME: Nω-Nitro-L-arginine methyl ester
NO: Nitric oxide
NOS: Nitric oxide synthase
NOX: NADPH oxidase
NP: Normotensive pregnant women
PE: Preeclampsia
qRT-PCR: Quantitative real-time polymerase chain reaction
RFU: Relative fluorescence units
RLU: Relative luminescence units
ROS: Reactive oxygen species
SBP: Systolic blood pressure
SEM: Standard error of the mean
siRNA: Small interfering RNA
SNP: Sodium nitroprusside
SOD1: Superoxide dismutase 1
TNF-α: Tumor necrosis factor alpha
U46619: Thromboxane A_2_ analog U46619

## WHAT IS NEW?

- Identifies NOX5 as a novel mediator of fetal vascular dysfunction in PE.
- Shows that preeclamptic umbilical arteries exhibit oxidative stress, reduced NO bioavailability, and selective NOX5 upregulation.
- Demonstrates that circulating factors from preeclamptic pregnancies transfer the hypercontractile phenotype to healthy umbilical arteries through a TNF-α/Ca²⁺/NOX5-dependent mechanism.
- Establishes that pharmacological inhibition or genetic silencing of NOX5 prevents oxidative stress and restores endothelial function.

## WHAT IS RELEVANT?

- Shows that PE directly impairs fetal vascular health, extending beyond the maternal and placental compartments.
- Uncovers the TNF-α/Ca²⁺/NOX5 axis as a mechanistic link between systemic inflammation, redox imbalance, and umbilical artery dysfunction.
- Positions NOX5 as a promising therapeutic target to preserve fetal vascular homeostasis in hypertensive pregnancies.

## CLINICAL/PATHOPHYSIOLOGICAL IMPLICATIONS?

- Suggests that targeting NOX5-dependent redox signaling may offer a strategy to mitigate pregnancy-related endothelial dysfunction.
- Indicates that part of nifedipine’s vascular benefit in PE may involve inhibition of Ca²⁺-dependent NOX5 activation.
- Highlights the possibility that preventing fetal vascular injury may have implications for long-term cardiovascular health.

